# Genomic epidemiology and multilevel genome typing of *Bordetella pertussis*

**DOI:** 10.1101/2023.04.26.538362

**Authors:** Michael Payne, Zheng Xu, Dalong Hu, Sandeep Kaur, Sophie Octavia, Vitali Sintchenko, Ruiting Lan

## Abstract

*Bordetella pertussis* is responsible for the respiratory infectious disease pertussis (or whooping cough), which causes one of the most severe diseases in infants, although it can be prevented by whole cell and acellular vaccines. The recent resurgence of pertussis is partially due to pathogen adaptation to vaccines as well as resistance to antimicrobials. Surveillance of current circulating and emerging strains is therefore vital to understand the risks they pose to public health. Although there is increased genomics based typing, a genomic nomenclature for this pathogen has not been well established. Here, we implemented the Multilevel Genome Typing (MGT) system for *B. pertussis* with five levels of resolution, which provide targeted typing of relevant lineages as well as discrimination of closely related strains at the finest scale. The low resolution levels can describe the distribution of alleles of major vaccine antigen genes such as *ptxP, fim3, fhaB* and *prn* as well as temporal and spatial trends within the *B. pertussis* global population. Mid-resolution levels enables typing of antibiotic resistant lineages and Prn deficient lineages within the *ptxP3* clade. High resolution levels can capture small-scale epidemiology such as local transmission events and has comparable resolution to existing genomic methods of strain relatedness assessment. The scheme offers stable MGT type assignments aiding harmonisation of typing and communication between laboratories. The scheme is available at www.mgtdb.unsw.edu.au/pertussis/ is regularly updated from global data repositories and accepts public data submissions. The MGT scheme provides a comprehensive, robust, and scalable system for global surveillance of *B. pertussis*.

## Introduction

*Bordetella pertussis* causes the respiratory infectious disease, pertussis (whooping cough) which is particularly severe in infants. Globally, 24 million cases and 160,700 deaths were estimated to have occurred in 2014 [1]. A whole cell vaccine (WCV) was introduced in the late 1940s to early 1950s which significantly reduced disease incidence and mortality [2,3]. Subsequently, an acellular vaccine (ACV) was introduced in the 1990s due to the side effects of the WCV. However, re-emergence of pertussis has been observed in the last two decades in countries with high vaccine uptake.

Studies have suggested that the selection pressure from vaccine-induced immunity has acted on *B. pertussis* evolution in recent years [4–7], and therefore the use of ACV may have been a significant contributor to the resurgence of pertussis [8]. Increased variation of ACV- antigen genes, a mismatch of alleles between the vaccine strain and the current circulating *B. pertussis* strains and the recent increase in Prn-negative isolates all point to vaccine selection [4-7,9–11].

*B. pertussis* may also be under selective pressure from macrolide antibiotics used for treatment of pertussis such as erythromycin or azithromycin [12]. Macrolide resistant strains of *B. pertussis* have recently emerged from the genetic background of ptxP1 strains in China [13–15] and very recently from ptxP3 background [16]. Although only a small number of macrolide resistant *B. pertussis* isolates have been reported in other countries, monitoring of the international spread of macrolide-resistant strains is of paramount importance [15].

Molecular techniques previously used for public health surveillance of *B. pertussis* each had shortcomings. Pulsed-field gel electrophoresis (PFGE) is difficult to compare between studies while multi-locus variable-number tandem-repeat analysis (MLVA) and traditional 7 gene multi-locus sequence typing (MLST) both lack resolution in *B. pertussis* [17]. Allele typing of vaccine antigen genes and surface protein genes offered improved resolution and is widely used [18–21]. However, all of these methods lack the resolution for fine scale typing required for detailed epidemiology in *B. pertussis* due to its very low genetic diversity.

A genotyping scheme based on single nucleotide polymorphisms (SNPs) divided *B. pertussis* into 42 SNP profiles (SPs) and six SNP clusters [4]. Subsequently, six epidemic lineages (ELs) were defined to describe local epidemic population structures within Australian SNP Cluster I isolates [22–24]. Although these typing method provided a granular representation of the *B. pertussis* population, they required manual curation and construction of phylogenetic trees for the description of the epidemic lineages.

Recently core and whole genome MLST schemes have been developed for *B. pertussis* [25–27]. These schemes provide high resolution typing, with the majority of isolates assigned a unique identifier. Additional analyses are then required to group these identifiers together to describe larger trends, which add complexity and require technical expertise. We previously developed a genome typing method called multilevel genome typing (MGT) and have applied it to several bacterial species of public health importance [28–31]. MGT consists of a series of MLST schemes (or levels) with increasing numbers of loci. Each isolate is assigned a sequence type (ST) at each level providing a simple nomenclature with a gradient of resolutions. In this study we developed an MGT scheme and an online database for *B. pertussis* and demonstrated its usefulness in describing known and novel population structures and epidemiological trends.

## Methods

### Genomes used in this study

The initial set of genomes used to create the MGT was made up of 4,797 genomes. Of these, 4,610 genomes passed MGT locus calling thresholds and were used to develop the MGT scheme [30]. These included 733 complete assemblies and 3,877 sets of paired-end Illumina reads collected from the NCBI RefSeq database and sequence read archive (SRA), respectively. A large number of isolates with paired-end Illumina reads were released after the MGT scheme was generated. These 1,930 sets of paired-end Illumina reads were added to previous datasets for a total of 6,540 isolates.

### Core genome definition including partial locus selection

Partial locus selection involved the division of reference loci in order to increase the proportion of isolates in which a locus can be reliably typed (typeability) and was composed of two steps. Firstly, all coding regions and intergenic regions from the *B. pertussis* Tohama I reference genome (GCF_000195715.1) with IS removed were initially selected and alleles were called as described previously (Supplementray methods A) [30]. Secondly loci with regions that were poorly typable were split on either side of that region (supplementary methods B). If the resulting sections of the locus (called subloci) were greater than 100bp, they were included as two or more separate loci in the core genome.

### MGT level locus selection

The low diversity and high clonality of *B. pertussis* results in many loci with only 2 alleles across the entire species. These loci are ideal for MGT definition because, when combined into one MGT level, they will not produce multiple minor STs caused by rare alleles in a small number of isolates. To best describe a clade in the overall population a locus should have as close to 2 alleles as possible, one for the clade of interest and another for the remainder of the population (supplementary figure 1). These loci were termed ‘phylogenetically informative loci’ and made up MGT levels 2, 3 and 4. Combining one locus for each target clade into one MGT level ensures that each clade should have a distinct ST with minimal small STs introduced by sporadic mutations. Additional level four loci were selected if they had at least 2 alleles, and all alleles were assigned to a minimum of 5 genomes. Finally, for a locus to be included an allele must be callable in all but 10 genomes (of 4,610) and must be partially callable in all but 20 genomes. In total an allele must be called in 4580/4610 genomes for a locus to be included in the MGT scheme.

### MGT assignment

MGT assignment used pipelines described previously and is hosted at the MGTdb [30,31] (supplementary methods C).

### Phylogenetic analyses

The allele profiles generated for each isolate that was successfully assigned an ST in the cgMLST scheme (n=4,610) were applied to produce a phylogeny of the whole dataset using GrapeTree v1.5.0, in which the rapidNJ algorithm was used [32]. A phylogeny composed of a single representative for each of the 82 MGT4 STs assigned to more than 5 genomes was used to simplify visualisation of MGT levels and other typing systems. The phylogeny was generated by extracting SNPs from each MGT5 allele relative to the reference allele (allele 1) for each isolate. SNPs were then concatenated into an alignment that was used to generate a maximum likelihood phylogeny using IQ-TREE v2.2.0 with default settings, 1,000 bootstrap pseudoreplicates and the K3Pu+F+I model automatically selected as the best-fit [33].

### prn gene mutation detection

Disruptions of the *prn* gene including SNPs, insertions, deletions and IS disruptions were all examined using a combination of methods and were compared to previously described events (supplementary methods F).

## Results

### B. pertussis core genome definition

Using 4,610 genomes, 4,633 core loci were identified with a total nucleotide length of 3.49 Mbp (85.12% of the Tohama I reference genome) which included 2,825 intact genes, 673 gene subloci, 860 intergenic regions and 275 intergenic region subloci (supplementary table 1). This core genome is much larger than the existing cgMLST scheme (1.75 Mbp) and comparable to the existing wgMLST scheme (3.34 Mbp) [25,27].

To facilitate locus selection for the MGT scheme levels, alleles were called for all core loci in the initial set of 4,797 isolates using standard MGT scripts. A minimum threshold of 96% of core loci must be successfully assigned an allele in order for an ST to be assigned to a genome [30]. This threshold was reached in 4,610 isolates and these were used in the further development of the MGT scheme. The average number of missing loci per genome is also an important measure of the typeability of the core genome. The core loci defined here had an average of 12.21 loci missing per isolate for 4,797 isolates. We compare that with the existing cgMLST scheme which had an average of 14.4 loci missing per isolate for 1,949 isolates [25]. Thus, the core loci allele typing performed here was reliable whilst providing significantly better resolution (Supplementary materials A).

### B. pertussis MGT scheme design

Due to the low genomic diversity observed in current *B. pertussis* isolates, five MGT levels were implemented rather than the eight or nine levels as in other species [28-30]. The first level is the well-established seven gene MLST [34]. MGT2, MGT3 and MGT4 loci were selected to best describe previously defined genotypes, clusters and lineages [4,22-24] (Figure 1). MGT4 loci were also partly selected to minimise small STs (assigned to five or fewer isolates). MGT5 loci were identical to the entire set of core genome loci defined above. Characteristics of each level are presented in Table 1.

**Figure 1.**
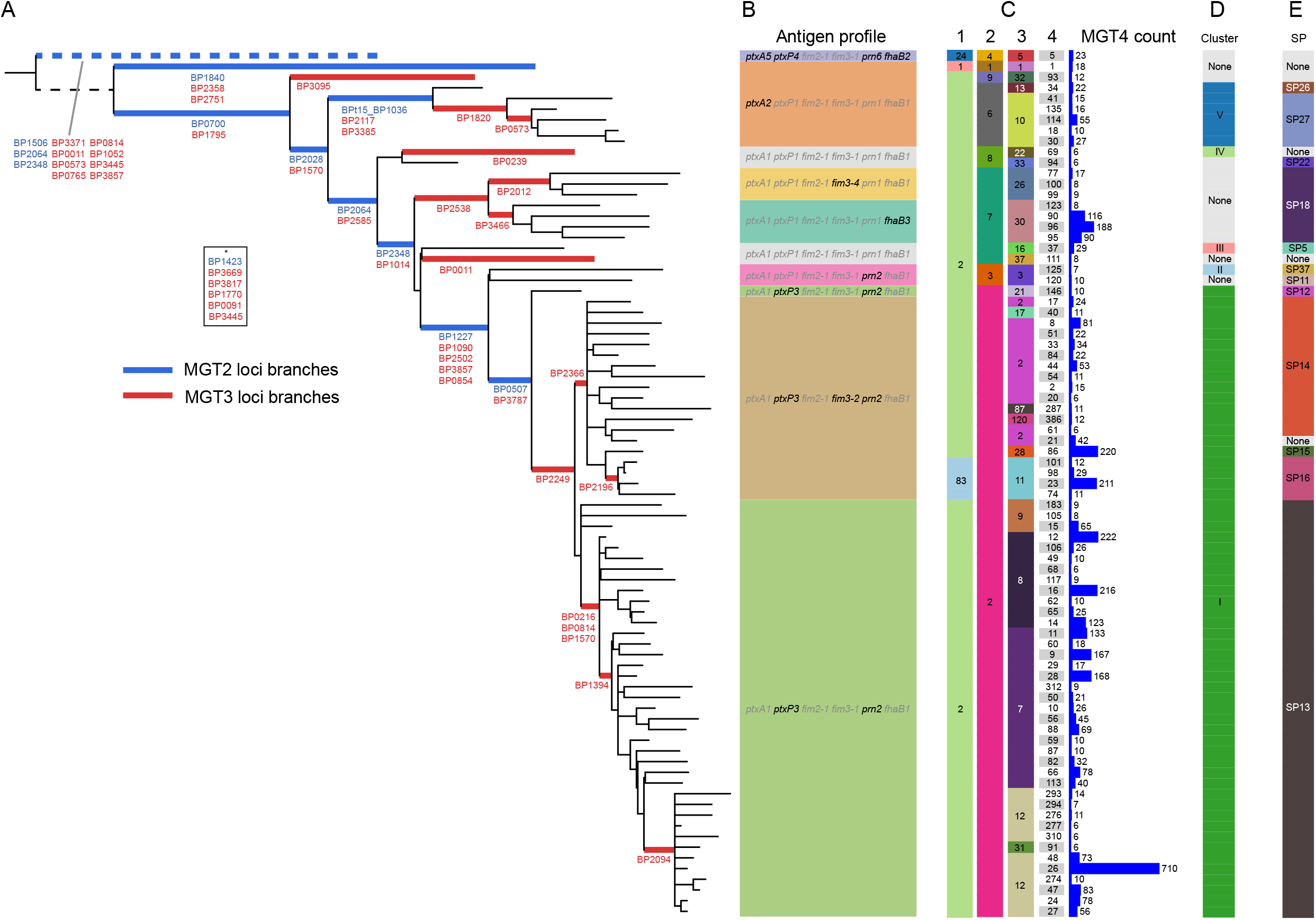
Representative phylogeny of MGT4 sequence types describing construction and correlation MGT2 and 3 with previous typing systems. **A**. A maximum likelihood phylogeny with each leaf representing every MGT4 ST with more than 5 genomes assigned to it. The position of allelic changes in loci that make up MGT2 and 3 are indicated by blue and red highlighting with the locus names presented below the relevant branch. Loci in box indicated by ^*^ are found on branches not present in this tree due to the 5 genome limit. **B**. MGT1, 2, 3 and 4 ST assignments for each leaf as well as the counts of each MGT4 ST in the 4610 genomes used in the MGT (435 genomes were assigned to MGT4 STs with less than 6 isolates and are not represented). C. Alleles for 5 common acellular vaccine antigens and ptxP for each MGT4 representative. D. Cluster and SNP profile assignments for each MGT4 ST.

**Table 1.** Characteristics of the 5 MGT levels.

| Level | Number of Loci | Nucleotides (% of Tohama I reference) | Number of STs assigned | Mean ST size | Number of isolates with failed ST (n=4797) | Avg failed allele calls per isolate | Avg partial allele calls per isolate |
| --- | --- | --- | --- | --- | --- | --- | --- |
| MGT1 | 7 | 2924 (0.07) | 4 | 1183 | 235 | N/A | N/A |
| MGT2 | 10 | 8820 (0.2) | 20 | 217.6 | 0 | 0.001 | 0.002 |
| MGT3 | 46 | 43653 (1.1) | 110 | 42.4 | 1 | 0.067 | 0.057 |
| MGT4 | 166 | 141212 (3.4) | 442 | 10.6 | 30 | 0.322 | 0.435 |
| MGT5 | 4633 | 3492117 (85.1) | 2772 | 1.8 | 192 | 12.21 | 26.45 |

### MGT STs describe previously established typing schemes

To allow comparison to existing studies the degree of agreement between existing typing systems and MGT STs was compared. MGT STs from MGT2 and MGT3 were able to describe alleles in the ACV antigen genes (*ptxA, fim2, fim3, prn, fhaB*) and the pertussis toxin promoter (*ptxP*) with 100% specificity and over 90% sensitivity (supplementary results B and C). Three SNP clusters and the majority of SPs defined previously [4] were well described by MGT2 and MGT3 STs (Figure 1D and E), while for the 6 Australian ELs [22-24], MGT4 STs provided highly specific typing (supplementary results D).

### B. pertussis global population structure described using the multiple levels of MGT

An updated global dataset of 6540 isolates (including the MGT definition dataset of 4610) was used to examine the genetic and geographical distributions of *B. pertussis* populations using MGT (Supplementary table 3). The most common MGT2 ST was ST2 (*ptxP3* allele), making up 82.3% (5382/6540) of the entire dataset followed by ST7 and ST6 with 9.8% (644/6540) and 3.2% (207/6540), respectively.

When applying the increased resolution of MGT3, 24 STs contained more than 10 isolates each and described 95.9% of the total sampled population (6274/6540, coloured nodes in Figure 2A). Importantly for the usefulness of these subdivisions, the large MGT2 ST2 was separated into 13 MGT3 STs, with the largest of these (ST7) making up 23.9% (1565/6540) of the total dataset. Interestingly, 98.0% (1166/1190) of all MGT3 ST12 genomes were found in North America while 98.5% (469/476) of MGT3 ST30 isolates were from China.

**Figure 2.**
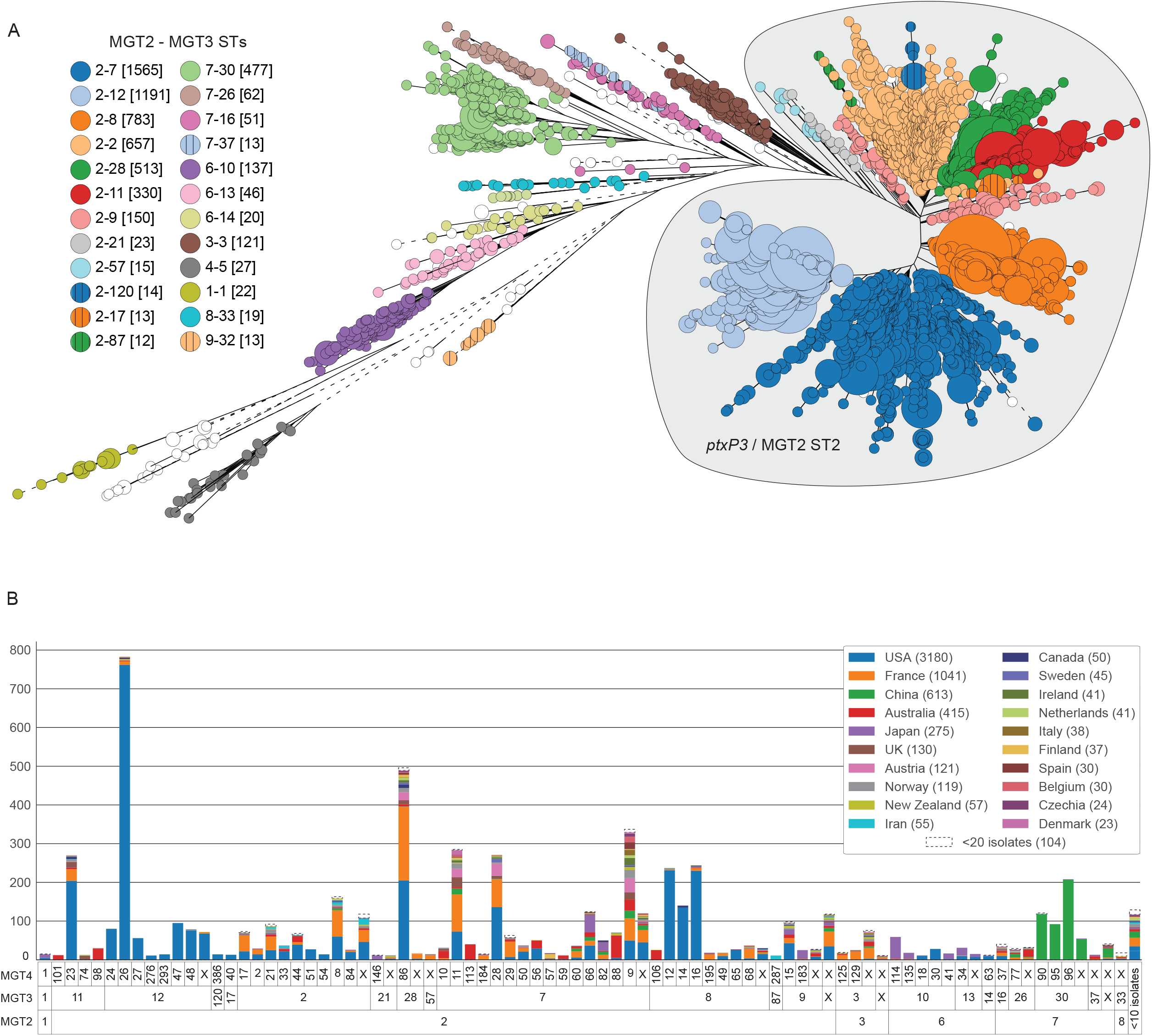
Global description of *B. pertussis* population structure using MGT. **A**. Evolutionary relationships between MGT2 and MGT3 STs. A neighbour joining tree constructed using rapidNJ within grapetree using MGT5 allele profiles from 6540 isolates. Each node represents one MGT5 ST with the size proportional to the number of isolates assigned. Nodes are coloured with MGT2 and MGT3 ST. The MGT2 ST2 clade (ptxP3) is shaded in grey. **B**. Each column is a unique set of MGT2, MGT3 and MGT4 STs with 10 or more isolates assigned to it. Colours are country of isolation within each ST set. Countries with less than 20 isolates are grouped together.

By further subdividing the population with MGT4 STs, more detailed country associations were identified (Figure 2B). Many MGT4 STs contained isolates from multiple continents, such as ST8, ST9 and ST86. However, a number of STs were more strongly associated with single countries (over 90% of samples from the same country). For example, in the USA, this included multiple MGT3 ST12 subtypes as mentioned above, but also three MGT3 ST8 subtypes, MGT4 ST12, ST14 and ST16. Other country associated STs included three in Japan (MGT4 ST183, ST146 and ST114), three in China (MGT4 ST90, ST95 and ST96) and four in Australia (MGT4 ST98, ST10, ST113 and ST106). Although France had the second highest number of sampled isolates (1041), there were no MGT4 STs uniquely associated with that country.

### Exploring population structure changes within the ptxP3 clade over time using MGT

In addition to establishing the spatial population differences, we also examined how the levels of MGT can describe temporal changes in population structure using ptxP3 strains (MGT2 ST2) [5,9,35]. MGT2 ST2 increased globally over time, making up 24.8% (99/399) of isolates before 2000, 78.4% (561/716) in the 2000s and 88.2% (4651/5274) in the 2010s. By using MGT3, the vast majority of MGT2 ST2 isolates (97.8%, 5263/5382) were assigned to 12 MGT3 STs (Figure 2A). The temporal changes of MGT3 STs are shown in Figure 3 and supplementary Figure 2. The two countries with the most sampled isolates were the USA (figure 3A) and France (figure 3B). These countries both showed a large change of population structure between 2008 and 2012 with MGT3 ST2 and ST9 being replaced by MGT3 ST7, ST8, ST11, ST12 and ST28. Interestingly, the dominant STs in each country during this change are different. In the USA, MGT3 ST12 was the most common followed by MGT3 ST8 and ST7 with MGT3 ST28 only becoming more common after 2015. In France MGT3 ST7, and subsequently MGT3 ST28, were the major drivers of population change with MGT3 ST8, ST11 and ST12 playing relatively minor roles. In the rest of the world the transition timeframe from old to new STs was the same and, as was observed in France, MGT3 ST7 was the dominant ST from 2009 onwards with smaller contributions from MGT3 ST11 and ST28 with little MGT3 ST8 or ST12 observed (Figure 3C).

**Figure 3.**
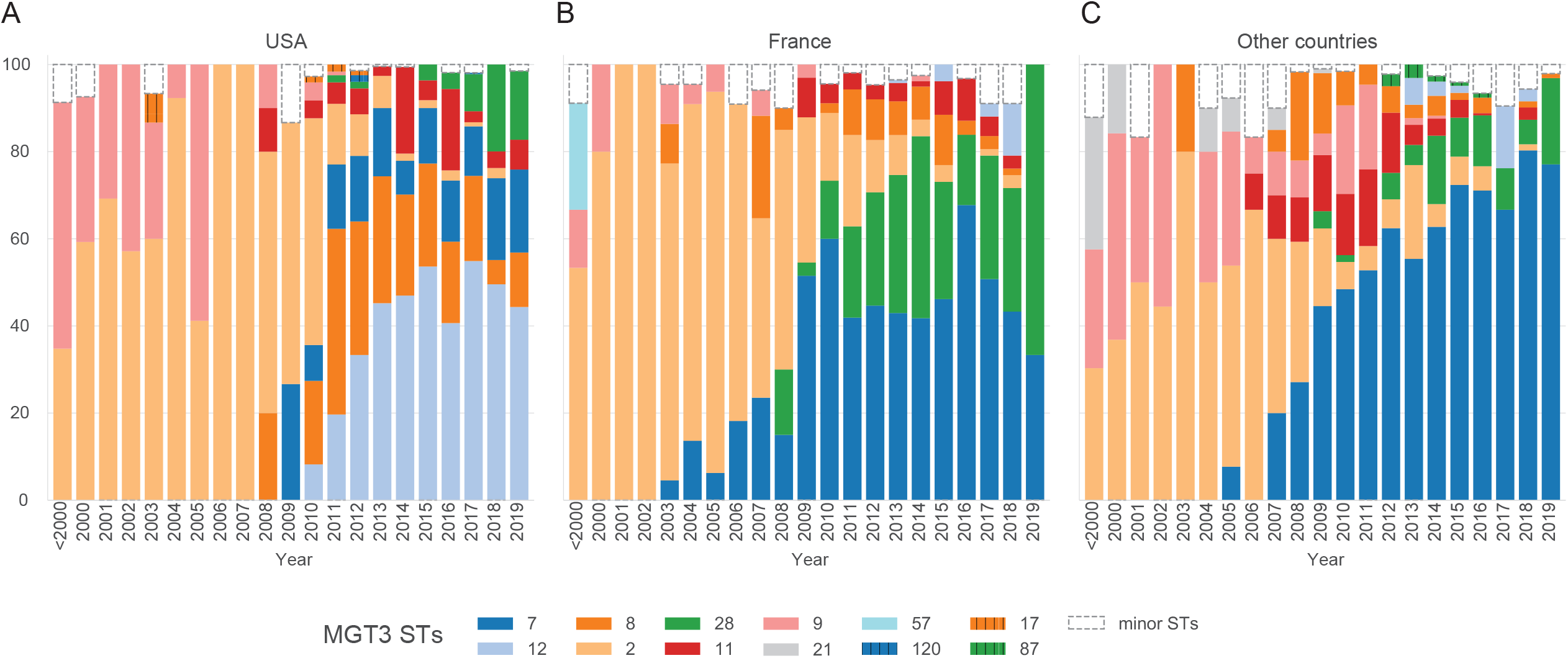
Temporal changes in MGT3 ST proportions describe evolution within the *ptxP3* (MGT2 ST2) clade. The proportion of isolates assigned to each major ST within MGT2 ST2 in each year are shown by different colours for the USA (**A**), France (**B**) and the rest of the world (**C**). Isolates assigned to minor STs are grouped together.

### Tracking macrolide resistant strains in China and globally

Macrolide resistant isolates were identified by typing the resistance conferring mutation, A2047G, in the 23S rRNA genes. Of 613 isolates from China, 488 (79.6%) carried the mutation. By MGT, MGT3 ST30 covered 76.1% (469/613) of the Chinese isolates and 93.4% (456/488) of Chinese macrolide resistant isolates. Using MGT4 STs, we can describe the previously reported macrolide resistant lineages 1, 2 and 3 as ST95, ST96 and ST90, respectively (Supplementary figure 3). In all, these three STs captured 84.8% (414/488) of resistant isolates in China. There were 15 isolates from outside of China predicted to be resistant which were found in 6 MGT4 STs and four countries (Supplementary table 4, supplementary results E). Importantly two isolates from Taiwan and one isolate from Japan belong to Chinese resistant lineages (MGT4 ST90 and MGT4 ST96 respectively).

### Multiple independent Prn deficient lineages described by MGT

The rapid increase of Prn deficient isolates (Prn-) in the last 15 years has been associated with multiple independent mutational events across the *B. pertussis* population [6,36]. Using known mutations that silence Prn production, we examined all 6540 isolates and identified 2963 isolates (45.3%) as Prn-. These isolates were found in 15 of 33 major MGT3 STs (2919 Prn- isolates) and 59 of 88 major MGT4 STs (2857 Prn- isolates) (Figure 4). Prn deficiency in these isolates was caused by 24 previously described mutations, 21 of which were previously confirmed by the Western Blot method (Supplementary tables 3 and 5, Supplementary figure 4). Of these mutations, 13 were assigned to more than 10 isolates (2833 isolates total) and 11 were assigned to 10 or fewer isolates (34 isolates total). An additional 200 isolates contained mutations whose effect on Prn production was unknown (marked uncertain).

**Figure 4.**
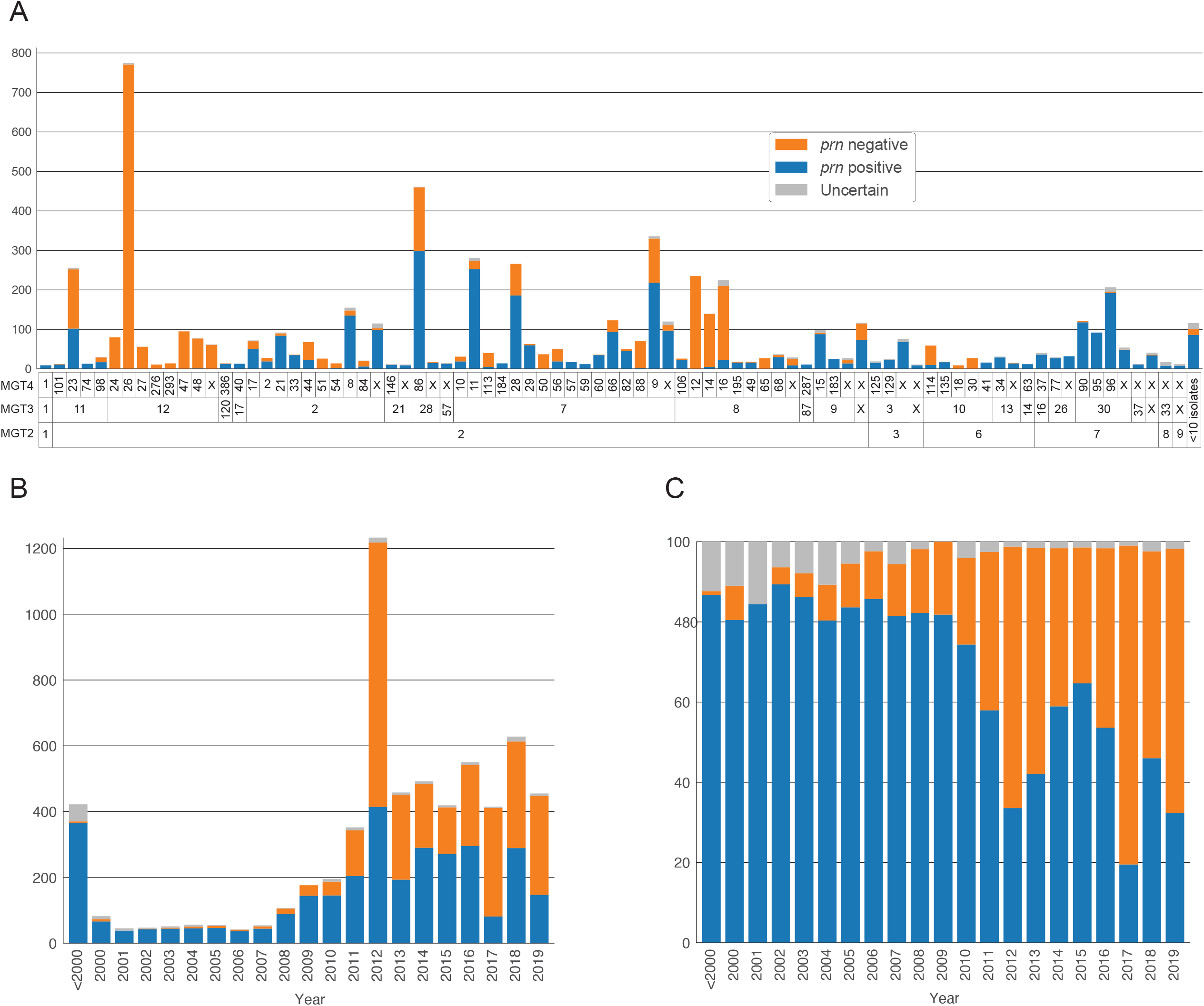
Distributions of *prn* negative isolates across both MGT types and years. **A**.Each column is unique set of MGT2, MGT3 and MGT4 STs with 10 or more isolates assigned to it. Isolates in each set are coloured by prn status. **B** and **C**. The absolute numbers and proportions of isolates that are prn negative, prn positive or uncertain in each year. In all plots orange is Prn negative, blue is Prn positive and grey is uncertain.

Prn- isolates are in the majority in many MGT3 and MGT4 STs. At MGT3 the two previously described USA dominated lineages (MGT3 ST12 and MGT3 ST8) were almost entirely Prn-, however these deficiencies were not caused by the same mutation. In MGT3 ST12, 99.92% (1190/1191) were Prn- with 99.66% (1187/1191) being caused by event 24 (an IS insertion at position 1608). Within MGT3 ST8 three large MGT4 STs were all majority Prn- but these deficiencies were caused by separate mutation events, for MGT4 ST12 (100% Prn-) event 12 (a nonsense mutation at position 1273), for MGT4 ST14 (96.53% Prn-) event 24 and for MGT4 ST16 (90.20% Prn-) event 56 (an IS insertion and deletion at position 240). MGT STs can therefore describe lineages that were Prn- and also distinguish between lineages with different causative mutations.

### MGT describes Australian epidemic lineages

At MGT2 level, Australian *B. pertussis* generally mirrored the trends seen globally with MGT2 ST2 making up 93.8% (225/240) of isolates sequenced since 2010. Similarly, at MGT3, the globally prevalent MGT3 ST7 accounted for 65.83% (158/240) of isolates since 2010 with MGT3 ST11, ST2 and ST8 accounting for 11.68%, 5.42% and 4.58% of isolates, respectively (Figure 5A). MGT3 ST12, which was predominant in the USA was only found twice in Australia.

**Figure 5.**
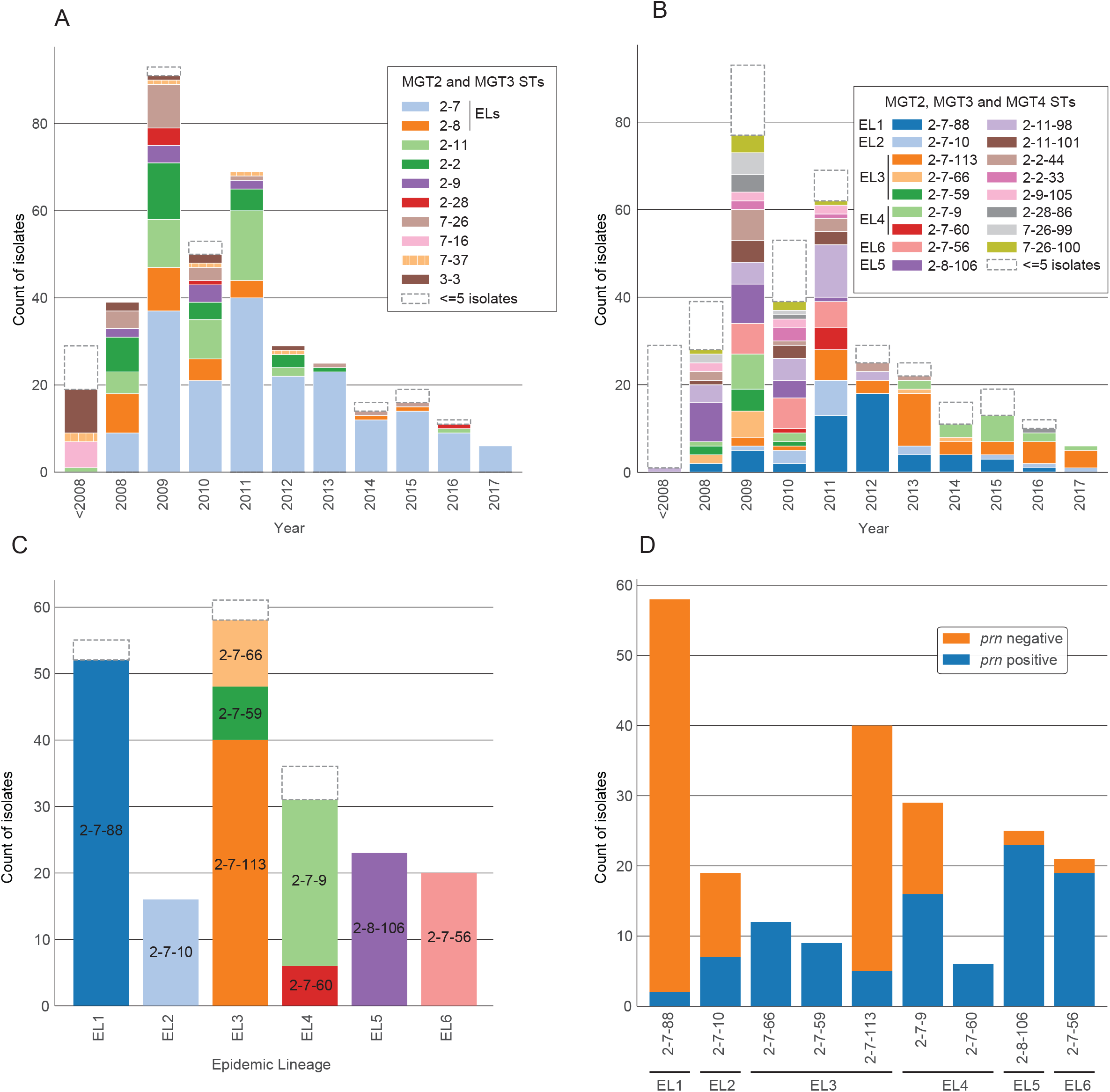
Description of all 390 Australian isolates using MGT. **A**. ST assignments per year using MGT2 and 3. STs containing the 6 epidemic lineages (ELs) are indicated. **B**. ST assignments per year using MGT2, 3 and 4. STs found in the six ELs are listed. **C**. The MGT2, MGT3 and MGT4 STs assigned to isolates in each EL. **D**. The prn status of Australian isolates assigned to major STs found in ELs. In all panels white dashed outline boxes represent the number of isolates assigned to STs with 5 or fewer isolates.

The six previously defined Australian epidemic lineages [23,24] can be described using MGT4 STs (Figure 5B). EL1, EL2, EL5 and EL6 were assigned to MGT4 ST88, ST10, ST106 and ST56 respectively. EL3 and EL4 were assigned to 3 and 2 MGT4 STs respectively, all of which were found within MGT3 ST7. The division of EL3 and EL4 into multiple STs was expected as these ELs contained more than one independent sublineage [22].

MGT4 allows the description of temporal patterns of the ELs as well as describing diversity within EL3 and EL4 (Figure 5C). EL5 and EL6 STs were not observed after 2011 while the remaining ELs were observed in most years. Within EL3, MGT4 ST59 and ST66 were only found in 2008, 2009 and 2010 while MGT4 ST113 emerged in 2009 and became the dominant ST in that EL and was identified in all subsequent years. In EL4, MGT4 ST9 was dominant and occurred in all years except 2011 and 2012 with the minor MGT4 ST60 only occurring in 2010 and 2011. Of the major STs associated with ELs 4 of 9 had significant numbers of Prn deficient isolates (MGT4 ST88, ST10, ST113 and ST9, figure 5D). These STs were also the only major STs observed after 2011 making up 82 of 110 isolates (74.55%), 80 of which were Prn deficient. Of minor STs in the same period 19/28 were also Prn deficient.

### MGT5 provides high resolution typing for local scale epidemiology

MGT5 provides high resolution typing for discriminating closely related isolates, which is vital given the low diversity of the *B. pertussis* population. Of the 6540 isolates in the global dataset, 3580 (54.74%) were assigned to singleton MGT5 STs while the remaining 2960 isolates were assigned to 720 MGT5 STs, each containing 2 to 75 isolates. We therefore investigated whether these STs could be used to infer relationships on small temporal and spatial scales.

To determine whether isolates sharing an MGT5 ST were more closely distributed than other isolates, pairwise comparisons of Australian isolate locations and dates were performed (n=268). Previous work identified that isolates within 1km of each other and isolated within 18 months had increased relative risk of transmission [22]. Of isolate pairs with the same MGT5 ST, 46.10% (284/616 pairs) were isolated within 1km of each other and within 18 months of each other. By contrast for isolate pairs with different MGT5 STs, this same parameter was 0.20% (144/71208 pairs). Therefore, isolates that share MGT5 STs are far more likely to represent recent, local transmission events than those with different MGT5 STs. This trend is also found when examining the frequencies of temporal and spatial distributions for all pairs of isolates within the same ST (Supplementary figure 5A) in comparison to pairs of isolates with different STs (Supplementary figure 5B).

The performance of MGT5 relative to a previous cgMLST scheme was evaluated using 55 isolates examined in that study [25]. Of the 55 isolates examined in the previous cgMLST scheme 6 STs were defined with more than 1 isolate (supplementary table 8). Of these STs, MGT5 was able to provide additional differentiation in 3 STs. In two other cases MGT5 grouped 2 isolates together that were separate in the previous scheme.

### B. pertussis MGT web database features

The *B. pertussis* MGT scheme is publicly available at https://mgtdb.unsw.edu.au/pertussis and the database includes all isolates in this study. The database is also open to public data submission including raw reads and processed allele files and all publicly available *B. pertussis* isolates are continuously updated as they are released from SRA. For publicly available isolates and raw read submissions, MGT types, vaccine antigen alleles and macrolide resistance SNP detection are called and reported per isolate. For processed allele submission, only MGT types and vaccine antigen calls are performed. The MGT database has an array of data export and visualisation features [31] that can facilitate epidemiological surveillance and outbreak investigation.

## Discussion

This study presents a novel MGT scheme for *B. pertussis* capable of providing analyses ranging from species-wide population structure (MGT2) to high-resolution genotyping enabling cluster/outbreak investigations within one country (MGT5). These MGT sequence types offer a simple nomenclature for communication and standardisation allowing integration and description of previously disparate datasets and analyses. Importantly, the existing, widely used vaccine antigen gene allele nomenclature as well as other SNP based typing systems were generally consistent with the MGT, allowing backwards compatibility with non-genomic studies and surveillance.

Given the low genomic diversity of the *B. pertussis* population, it is essential to utilise the largest possible proportion of the genome to provide maximum discrimination between closely related isolates. MGT provides this high resolution as it utilised intergenic regions as well as fragments of genes to increase the overall resolution of the system. These steps mean that the MGT5 level (*B. pertussis* cgMLST) has equivalent resolution to an existing wgMLST scheme while maintaining the standardisation that is inherent in a cgMLST scheme [25,27]. The performance of MGT5 relative to the existing cgMLST scheme demonstrates that this increased resolution allowed closely related isolates to be further differentiated. The cases where an MGT5 ST was split into 2 cgMLST STs were likely due to differences in allele calling algorithms that lead to mutations in one scheme not being called in another. The performance of MGT5 typing in describing isolates that are closely related temporally and spatially suggest that they will be useful in describing small scale epidemiology.

As well as providing high resolution at MGT5 the lower levels of the MGT (MGT2, 3 and 4) have been designed to describe lineages of interest and to match the underlying genetic relationships of the population as closely as possible. The ability to generate definitive, stable sequence types at multiple resolutions provides an effective system for surveillance of strain movement nationally and internationally. This is especially the case for lineages that are not frequently observed in more than one location. For example, MGT3 ST30, and its three resistant lineages, MGT4 ST95, ST96 and ST90, were almost exclusively found in China [15]. Given that macrolides are the first line treatment for pertussis and also used as prophylaxis [37], the spread of macrolide resistant strains internationally could have significant global public health impacts. Effective surveillance for these lineages is therefore essential and MGT can perform this role without requiring reanalysis of the whole dataset as demonstrated by the description of small numbers of these lineages in Japan and Taiwan. Similarly, the MGT3 ST12 is almost entirely restricted to the USA where it is the most prevalent type. While any differences in the virulence and severity of disease caused by this clade have not yet been established, its dominance in the USA and potential international spread would also be of interest.

In all cases in this and any genomic epidemiology study, conclusions related to ST proportions and distributions are dependent on the current set of sequenced strains being representative of the larger population. The geographic restriction of MGT3 ST12 to the USA could be explained by lack of sampling in other locations. However, three other MGT3 STs (ST7, ST11 and ST28) emerged within the same 4-year period (2008-2011) and were commonly found in multiple continents, suggesting that MGT3 ST12 had not spread to other countries. The same was likely to be true for the Chinese restricted MGT3 ST30. These observations suggest that there are populations that are geographically restricted and maintained over at least 10 years in some locations. By contrast, no large geographically restricted STs were observed in France despite similar numbers of isolates being sampled. The reasons behind these differences are currently unclear.

The effective surveillance of *ptxP3* strains (MGT2 ST2) and their subtypes is vital given its global dominance. Existing nomenclatures, such as fim3 alleles or SNP based approaches, have been used to describe the evolution of different groups within *ptxP3* [4,6,22-24], however each has limitations. *fim3* alleles can only divide *ptxP3* into 2 types therefore providing only a marginal improvement in resolution. SNP based approaches work well, but they are static. New clades identified after the SNP-based approach’s development may not be described at all. In addition, they are currently not incorporated into a typing tools or maintained in a database. Consequently, they often require manual analysis and type assignment.

By contrast MGT typing can divide the *ptxP3* strains into numerous subtypes from MGT3 to MGT5, which enabled detailed description of its epidemiology. For example, *Prn* deficient isolates have been shown to have a fitness advantage [6,38] and have increased in frequency in both Australia and the USA. However, the genetic cause of the loss of prn production varies between these countries indicating that these strains were of independent origin. By applying MGT, it becomes clear that the increase in Prn deficient strains in the USA was caused by STs that are mostly distinct from those in France and the rest of the world.

We previously studied the molecular epidemiology of Australian *B. pertussis* using both gene typing and genome sequencing [22,24,35]. The strains responsible for the past 2 epidemics from 2008 to 2017 in Australia (grouped into epidemic lineages) can be clearly disentangled using STs at MGT3 and MGT4 levels. MGT allows the continued surveillance of these epidemic lineages. This ability to perform ongoing surveillance allowed the identification of 2008 to 2012 epidemic STs that are now likely extinct (e.g. MGT4 ST56 and ST106) and those that continued to cause infections in Australia (MGT4 ST88, ST10, ST113, and ST9). MGT also clearly demonstrates that the majority of isolates in STs after 2011 were Prn deficient, supporting previous work demonstrating the increased fitness of such strains [6,22,38].

In conclusion, this study developed an MGT system for *B. pertussis* and showed that different MGT levels provide a distinct and useful characterisation of the *B. pertussis* population. MGT level 2 can represent vaccine antigenic allele types, MGT level 3 can describe the evolution of the *ptxP3* strains, while MGT level 4 can identify independent Prn deficient lineages, lineages that caused the recent Australian epidemic and the three distinct Chinese erythromycin-resistant lineages. MGT level 5 offers the highest resolution for examining small scale temporal and spatial epidemiology. The MGT typing platform is publicly available and contains a continually updated database that provides visualisation, data export and typing to facilitate the adoption of this technology.

## Acknowledgements

The authors would like to acknowledge Robin Heron for technical assistance. This work was supported by The National Health and Medical Research Council: [Grant Number 1146938].

## Declaration of interests

The authors report there are no competing interests to declare.

## Data availability statement

All data used in this study are publicly available and all MGT data is available at mgtdb.unsw.edu.au/pertussis.

## Notes

https://mgtdb.unsw.edu.au/pertussis/

### Competing Interest Statement

The authors have declared no competing interest.

