## Supplementary Materials for "Genomic epidemiology and multilevel genome typing of *Bordetella pertussis*"

**Supplementary Methods**

A – Core genome allele calling

Alleles were called for all loci using the MGT reads2alleles script (github.com/LanLab/MGT_reads2alleles) and the typeability of the loci was examined. The MGT allele calling pipeline marked paralogous loci and loci with more than 20% of sequence missing as uncallable (0 allele). Additionally, loci that lack between 0 and 20% of sequence length are called with missing regions denoted by N and are defined as partially callable.

B – Core subloci definition

To identify regions of loci that were not reliably callable and should be excluded each locus typabiluty was examined for all isolates (n=4,797) and at each nucleotide position, a count of how many isolates were missing information was collected. A typability cut-off (5.5%) was used to determine what percentage of isolates a nucleotide position could be missing in before it was called as untypeable. The cut-off was selected that was the lowest possible while minimising the amount of the core genome that was removed. If a locus contained a region of nucleotides that exceeded this cut-off the locus would be split at either side of that region.

C- MGT assignment

Assemblies were generated using the shovill pipeline with skesa as the assembler (<http://github.com/tseemann/shovill>)[1]. Alleles were then assigned for each isolate and the combination of alleles was used to assign sequence types at each level. The combination of these sequence types is the genome type (GT). STs for MGT level 1 are assigned using mlst (<https://github.com/tseemann/mlst>) which employs the existing seven-gene MLST scheme [2]. Full pipeline description and further information is available at the MGT documentation (<https://mgt-docs.readthedocs.io/en/latest/analysis_pipeline.html>) [3].

*D - Core genome typability measurement (including IS disruption performance)*

The locations of insertion sequences in the complete genomes were determined using BLAST. Locations of cgMLST loci in complete genomes were determined using the MGT reads_to_alleles.py script (github.com/LanLab/MGT_reads2alleles). Where the IS location overlapped a locus location a locus was called as disrupted. For each IS insertion the isolate allele calls for both complete and illumina based assemblies were extracted from allele profiles to determine the accuracy of allele calls where the locus is disrupted by an IS. In order to compare the performance of the MGT pipeline to existing schemes, the typability of the existing cgMLST scheme was calculated by downloading all available allele profiles from the database website (https://bigsdb.pasteur.fr/bordetella/) and counting the number of loci called “N” for each isolate before the average across all isolates was taken.

*E - Vaccine antigen allele calling from MGT alleles*

*ptxA*, *ptxP*, *fim2* and *fim3* all used exact allele matches to identify MGT alleles corresponding to published allele sequences. Since accurately assembling the tandem repeat region in *prn* from short read data is extremely challenging, only SNPs were taken into account to differentiate alleles. Consequently, several alleles that are distinguished based on tandem repeat number variation were collapsed into assignments containing multiple allele numbers. For clarity *prn*1,12,7 is named *prn*1, *prn*2,3,4,5,9,13,15,17 is named *prn*2, *prn*6,10 is named *prn*6 and *prn*11,18 is named *prn*11. Due to the extremely large size of the *fhaB* gene, traditional allele typing has been limited to small subsections of the locus containing individual SNPs (Supplementary materials tables 1 and 2). Therefore, MGT alleles with SNPs corresponding to the *fhaB*1, 2 and 3 alleles were identified and assigned to one of the three allele types.

Supplementary materials table 1. SNPs at position 1334 and 4171 that define *fhaB* major typing alleles.

|  | fhaB subloci 2 positions | |
| --- | --- | --- |
| *fhaB* major allele | 1334 | 4171 |
| *fhaB1* | A | C |
| *fhaB2* | C | C |
| *fhaB3* | A | T |

Supplementary materials table 2. Alleles for the *fhaB* gene sublocus BP1879_sub_2 and corresponding *fhaB* typing SNP and major allele assignment.

| MGT allele | 1334 | 4171 | *fhaB*-major allele |
| --- | --- | --- | --- |
| 1 | A | C | 1 |
| 2 | C | C | 2 |
| 3 | C | C | 2 |
| 4 | A | C | 1 |
| 5 | A | C | 1 |
| 6 | A | C | 1 |
| 7 | A | C | 1 |
| 8 | A | T | 3 |
| 9 | A | C | 1 |
| 10 | C | C | 2 |
| 11 | A | C | 1 |
| 12 | A | C | 1 |
| 13 | A | C | 1 |
| 14 | A | C | 1 |
| 15 | A | T | 3 |
| 16 | A | T | 3 |
| 17 | A | C | 1 |
| 18 | A | T | 3 |
| 19 | A | T | 3 |
| 20 | A | T | 3 |
| 21 | A | C | 1 |

F – Prn disruption detection

Three different methods were used to detect Prn disruptions. Firstly, the *prn* coding sequence was used to search assembled and complete genomes using BLASTn (default parameters except min_raw_gapped_score was set to 100) in order to detect insertions and deletions. Secondly, for complete genomes all IS sequences in the ISfinder database were searched against each genome using BLASTn and cases where the IS was found within the *prn* gene were recorded [4]. Thirdly, snippy was used to identify point mutations and small indels that may disrupt *prn* production (<https://github.com/tseemann/snippy>). IS insertions were also putatively identified when the two following conditions were met. Firstly, the assembly of two parts of the *prn* gene were found at the ends of two contigs, and, secondly, the existence of a 6bp repeat (characteristic of some IS families) found at the end of each of those contigs. All mutations were compared to previous analyses and previously described events were noted [5-9].

**Supplementary Results**

*A - Core genome typing performance*

In *B. pertussis*, insertion sequences (IS) often disrupt loci leading to fragmented assembly and typing complications. The performance of short read allele calling and the influence of IS insertions can be assessed by comparing allele calls obtained from complete genomes to those obtained from assembled short reads of the same isolate. This comparison was performed on 559 complete genome/ illumina data pairs to evaluate MGT typing performance (supplementary methods C). 427 genomes had no mismatches between complete and short read assemblies while 91 had a mismatch at one locus, 27 at two loci and 10 at three or more loci. It is worth noting that these differences could be attributed to errors in the complete genome or the illumina assembly in the MGT pipeline, as well as the introduction of SNPs due to the use of different sequencing technologies or from passaging of strains between sequencing runs. In the 559 genomes compared, a total of 282 core loci were disrupted by IS insertion and of these 251 (89%) were called the same allele in both the complete and short read assemblies. The remaining 11% was made up of cases where an allele was not called in the complete and/or short read assembly. Hence, the MGT allele typer accurately manages IS insertions without generating erroneous allele calls. Overall, the core genome selected had good typability with an average of 12.21 loci missing and 26.45 loci partially callable per isolate.

*B – MGT2 and MGT3 STs correspond to major vaccine antigen alleles*

Alleles in the ACV antigen genes (*ptxA, fim2, fim3, prn, fhaB*) and the pertussis toxin promoter (*ptxP*) have been typed with multiple alleles. MGT describes each antigen allele with one or more MGT2 and MGT3 STs. The major alleles for each of the 6 antigen loci are found in monophyletic lineages. The agreement between these lineages and major MGT STs (assigned to more than 5 isolates) are listed in Supplementary materials table 3 and visualised in Figure 1 (all alleles and lineages are listed per isolate in Supplementary table 3).

For *ptxA, ptxP, fim2* and *prn,* MGT2 STs can differentiate major alleles while for *fim3* and *fhaB* MGT2 and MGT3 STs are required. When using MGT STs to infer antigen allelic types, all major and minor MGT STs are 100% specific to one antigen allele type except for one erroneous assignment of a *ptxP1* isolate to MGT2 ST2. MGT STs indicated in Supplementary Materials Table 3 are also at least 94% sensitive, meaning that at least 94% of isolates in an antigen allele type will have one of the indicated major STs with the remaining isolates all assigned to minor STs. Vaccine antigen alleles can also be accurately called directly from their loci, which are included in MGT level 5 (Supplementary results C).

Supplementary Materials Table 3. Performance of MGT2 and MGT3 STs in describing the same clades as vaccine antigen alleles

| **Antigen** | **Antigen allele** | **Specific MGT2 STs** | **Specific MGT3 STs** | **Proportion of antigen alleles described with specific MGT STs** | |
| --- | --- | --- | --- | --- | --- |
|  |  |  |  | **Percentage** | **Proportion** |
| *ptxA* | *ptxA1* | 2,3,7,8 | - | 99.74 | 4325/4336 |
|  | *ptxA2* | 1,6,9 | - | 97.29 | 215/221 |
|  | *ptxA5* | 4 | - | 100.00 | 27/27 |
| *ptxP* | *ptxP1* | 1,3,6,7,8,9 | - | 98.72 | 853/864 |
|  | *ptxP2* | 5 | - | 100.00 | 9/9 |
|  | *ptxP3* | 2 | - | 99.89 | 3693/3697 |
|  | *ptxP4* | 4 | - | 100.00 | 27/27 |
| *fim2* | *fim2-1* | 1,2,3,4,6,7,8,9 | - | 99.63 | 4582/4599 |
|  | *fim2-2* | 5 | - | 100.00 | 9/9 |
| *fim3* | *fim3-1* | - | all except below | 94.66 | 3441/3635 |
|  | *fim3-2* | - | 2,11,17,28,87,120 | 93.34 | 869/931 |
|  | *fim3-4* | - | 26 | 100.00 | 36/36 |
| *prn* | *prn1* | 1,5,6,7,8,9 | - | 98.54 | 808/820 |
|  | *prn2* | 2,3 | - | 99.87 | 3756/3761 |
|  | *prn6* | 4 | - | 100.00 | 27/27 |
| *fhaB* | *fhaB1* | 1,2,3,6,7**,8,9 | - | 99.71 | 4089/4101 |
|  | *fhaB2* | 4,5 | - | 100.00 | 36/36 |
|  | *fhaB3* | 7** | 30** | 93.02 | 440/473 |
| *in all cases isolates not in the correct MGT STs are in minor STs containing 5 or fewer isolates that are themselves specific to a single antigen allele  ** MGT2 ST7 contains both *fhaB2* and *fhaB3*. Within it at MGT3 level, only MGT3 ST30 is *fhaB3*. | | | | | |

*C - Antigen types can be directly called from MGT allele calls*

All 6 antigen/promoter loci (*ptxA, fim2, fim3, prn, fhaB*, *ptxP*) are found within MGT5 (the core genome defined in this study). It is therefore possible to call antigen types from allele calls produced during MGT processing. To obtain a baseline for the accuracy of these antigen types, lineages for each antigen allele were assigned based on the position of the relevant mutation in a whole dataset phylogeny generated using MGT5 allele profiles. Allele calls from MGT alleles were then compared to the antigen lineage of isolates to determine their accuracy (Supplementary materials table 4). All antigens genes, except for *fim3*, had novel identified alleles that have not been published before. In most cases, these were relatively uncommon, however, in *prn*, they were present in 6.3% of isolates with a single novel allele detected in 225 isolates. When the novel alleles are observed in multiple isolates they clustered together in phylogenetic clades (excluding *prn1005*), suggesting that they are unlikely to be assembly or sequencing errors (data not shown). When taking into account both the accurate calls and novel alleles, all 6 antigen loci were correctly assigned in over 99.5% of isolates. However, there were 30 isolates not able to be assigned one or more antigen alleles directly from allele calls. In these cases, a definitive MGT STs could be assigned as a proxy (supplementary materials table 5).

Supplementary materials Table 4. Performance of vaccine antigen allele calling from MGT allele calls. (N=4610)

| **Locus** | **Correct calls** | **Not called in MGT** | **Novel allele calls** | **Incorrect calls** | **% correct or novel** |
| --- | --- | --- | --- | --- | --- |
| *ptxA* | 4597 | 7 | 6 | 0 | 99.85 |
| *ptxP* | 4587 | 5 | 18 | 0 | 99.89 |
| *fim2* | 4601 | 1 | 1 | 7 | 99.83 |
| *fim3* | 4609 | 1 | 0 | 0 | 99.98 |
| *Prn* | 4295 | 14 | 295 | 6 | 99.57 |
| *fhaB* | 4605 | 3 | 1 | 1 | 99.91 |

Supplementary materials Table 5. Strains where MGT STs can be used to infer vaccine antigen allele.

| Strain | MGT2 | MGT3 | missing antigen allele | antigen allele inferred from MGT ST |
| --- | --- | --- | --- | --- |
| ERR029318 | 7 | 49 | fhaB | fhaB1 |
| L1191 | 3 | 35 | fhaB | fhaB1 |
| ERR037430 | 3 | 3 | fhaB | fhaB1 |
| SRR12105028 | 2 | 7 | fim2,prn | fim2-1,prn2 |
| GCA_002240515.1 | 2 | 7 | prn | prn2 |
| SRR8652768 | 2 | 7 | prn | prn2 |
| SRR5080692 | 2 | 7 | prn | prn2 |
| SRR11048617 | 2 | 7 | prn | prn2 |
| SRR12105146 | 2 | 7 | prn | prn2 |
| SRR12105049 | 2 | 7 | prn | prn2 |
| SRR12105086 | 2 | 7 | prn | prn2 |
| SRR12105120 | 2 | 7 | prn | prn2 |
| SRR12105119 | 2 | 7 | prn | prn2 |
| L2249 | 2 | 7 | prn | prn2 |
| SRR12105032 | 2 | 7 | prn | prn2 |
| SRR12105063 | 2 | 7 | prn | prn2 |
| L2244 | 2 | 7 | prn | prn2 |
| SRR8652757 | 2 | 7 | prn | prn2 |
| SRR9131490 | 12 | 50 | prn | prn2 |
| ERR029326 | 2 | 9 | fim3 | fim3-1 |
| GCA_001509915.1 | 2 | 8 | ptxA,ptxP | ptxA1,ptxP3 |
| GCA_002892705.1 | 2 | 8 | ptxA,ptxP | ptxA1,ptxP3 |
| SRR9002862 | 2 |  | ptxA,ptxP | ptxA1,ptxP3 |
| SRR9002901 | 2 | 22 | ptxA | ptxA1 |
| SRR9002822 | 2 | 12 | ptxA | ptxA1 |
| GCA_001509895.1 | 2 | 2 | ptxA,ptxP | ptxA1,ptxP3 |
| SRR9002815 | 2 | 28 | ptxP | ptxP3 |
| SRR10002425 | 2 |  | ptxA | ptxA1 |
| SRR9131393 | 2 | 2 | fhaBunknown | fhaB1 |

*D - Correspondence of* MGT with *previous SNP typing systems*

The relationship between SNP clusters/SPs and MGT STs from multiple levels was examined. The majority of the isolates (84.2%, 3884/4610) were assigned to a SNP cluster and a fair proportion of the unsigned isolates belonged to SP18 which is a known outlier outside of the SNP clusters. Six SNP clusters have previously been defined and were all detected in MGT data (supplementary table 6, Figure 1D). Cluster I corresponded to MGT2 ST2 which described 99.9% (3676/3680) of the cluster. Cluster V corresponded to MGT2 ST6 which described 99.4% (170/171) of the cluster. Cluster IV was described completely by MGT3 ST 22 (7 isolates). Clusters II and III were only found as a subset of single MGT3 STs (ST3 and ST16 respectively) while cluster VI was only identified in a single isolate with MGT3 ST111. Importantly, a total of 728 isolates (including all 445 isolates in MGT3 ST30) could not be assigned to any cluster. Additionally, Clusters II, III, IV and VI are nested within larger phylogenetic clades that are not precisely captured using the cluster-defining SNPs.

SPs were assigned to 94.6% (4363/4610) of isolates (Figure 1E) and SPs were matched to MGT2 and MGT3 STs. The relationships of SPs to MGT2 and MGT3 STs were broadly split into three categories. The first group included SPs (SP13,18 and 14) that described multiple MGT2 and MGT3 STs (7, 4 and 4 STs respectively). The second group included SPs (SPs 8,12,15,16,25,26 and 36) that correspond closely to specific MGT3 STs (ST51, ST21, ST28, ST11, ST14, ST13 and ST1 respectively). The third group consisted of the remaining 15 SPs, which were assigned to fewer than 35 isolates each, except for SP27 (which was found in 116 isolates). Multiple of these SPs were found within single MGT3 STs (See Supplementary table 7 and Figure 1.)

Six ELs were previously defined and assigned to 211 Australian isolates, all of which fell within MGT2 ST2 [10-12]. The MGT4 STs that were assigned to isolates in each EL were identified as detailed below. EL1,2,5 and 6 contained a single ST (MGT4 ST88, ST10, ST106 and ST56 respectively), EL4 contained two STs (MGT4 ST49 and ST60) and EL3 contained three STs (MGT4 ST59, ST66 and ST113).

E – Macrolide resistant isolates outside China

Of the 15 macrolide resistant isolates identified outside of China Four isolates from France have previously been described as resistant and belonged to MGT4 ST23 [13,14]. Of the 8 isolates from the USA, 5 belonged to MGT4 ST15 and have been described previously [15-17], two belonged to MGT4 ST37, one of which has been described previously [18] and one assigned MGT4 ST22 has not been described as resistant previously and occurred in the USA restricted MGT3 ST12 in 2017. Two isolates from Taiwan were within the MGT4 ST90 resistant lineage in 2011 and 2012 while one isolate from Japan was within the MGT4 ST96 resistant lineage in 2018.
