## Supplementary Figure 1 for "Genomic epidemiology and multilevel genome typing of *Bordetella pertussis*"

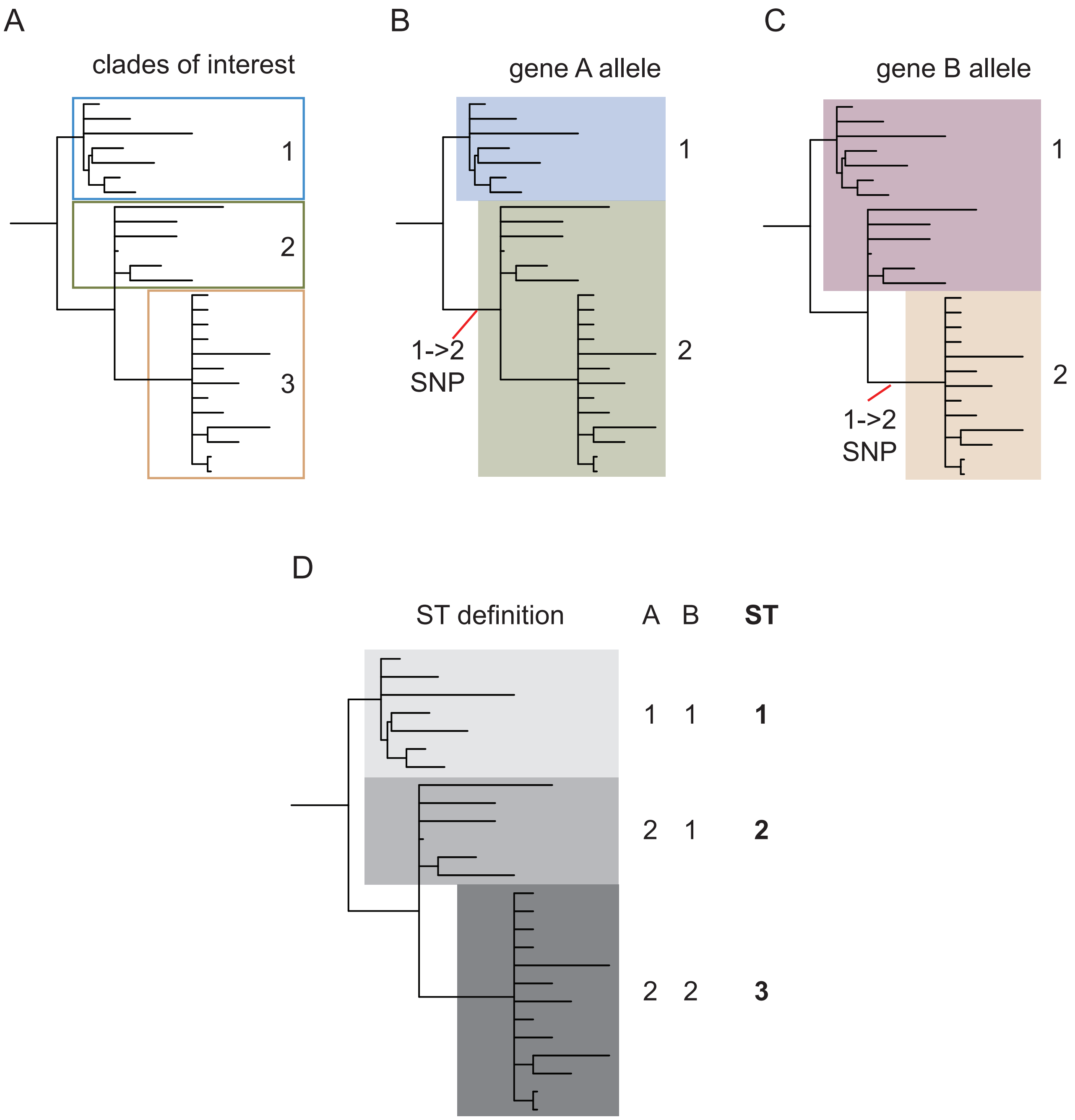


**Supplementary figure 1.** Phylogenetically informative loci selection for MGT2 and MGT3. **A.** An example phylogeny showing three clades of interest. **B.** Gene A has a SNP on the ancestral branch of clades 2 and 3 that leads to geneA having one allele for clade 1 and another for clades 2 and 3. **C.** A similar example to A with gene B alleles distinguishing clade 3 from clades 1 and 2. **D.** Combination of alleles in gene A and gene B into an allele profile defines three distinct STs that act as a nomenclature for the clades of interest.
