## Supplementary Figure 2 for "Genomic epidemiology and multilevel genome typing of *Bordetella pertussis*"

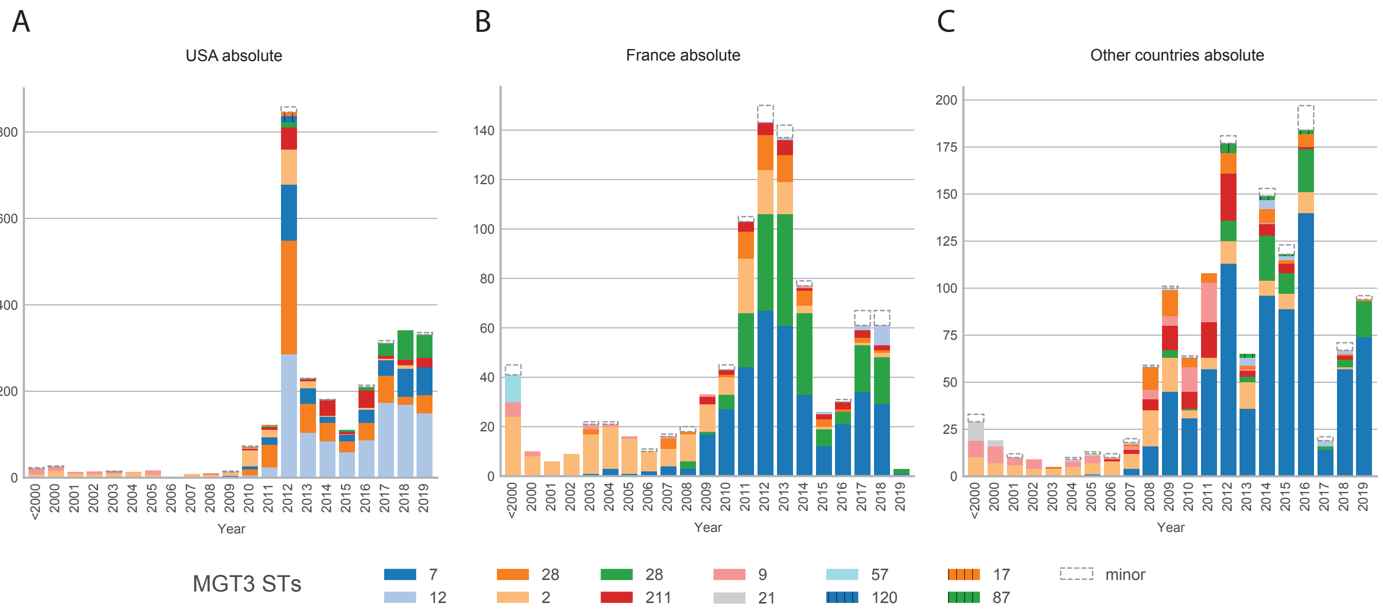


**Supplementary Figure 2. Temporal changes in MGT2-MGT3 ST counts.**The count of isolates assigned to each major MGT3 ST within MGT2 ST2 in each year are shown by different colours in each column. **A.** Counts of STs in the USA over time. **B.** Counts of STs in the France over time. **C.** Counts of STs in the countries other than the USA and France over time. Isolates assigned to minor STs within MGT2 ST2 are grouped together and are indicated by a dashed outline.
