## Supplementary Figure 3 for "Genomic epidemiology and multilevel genome typing of *Bordetella pertussis*"

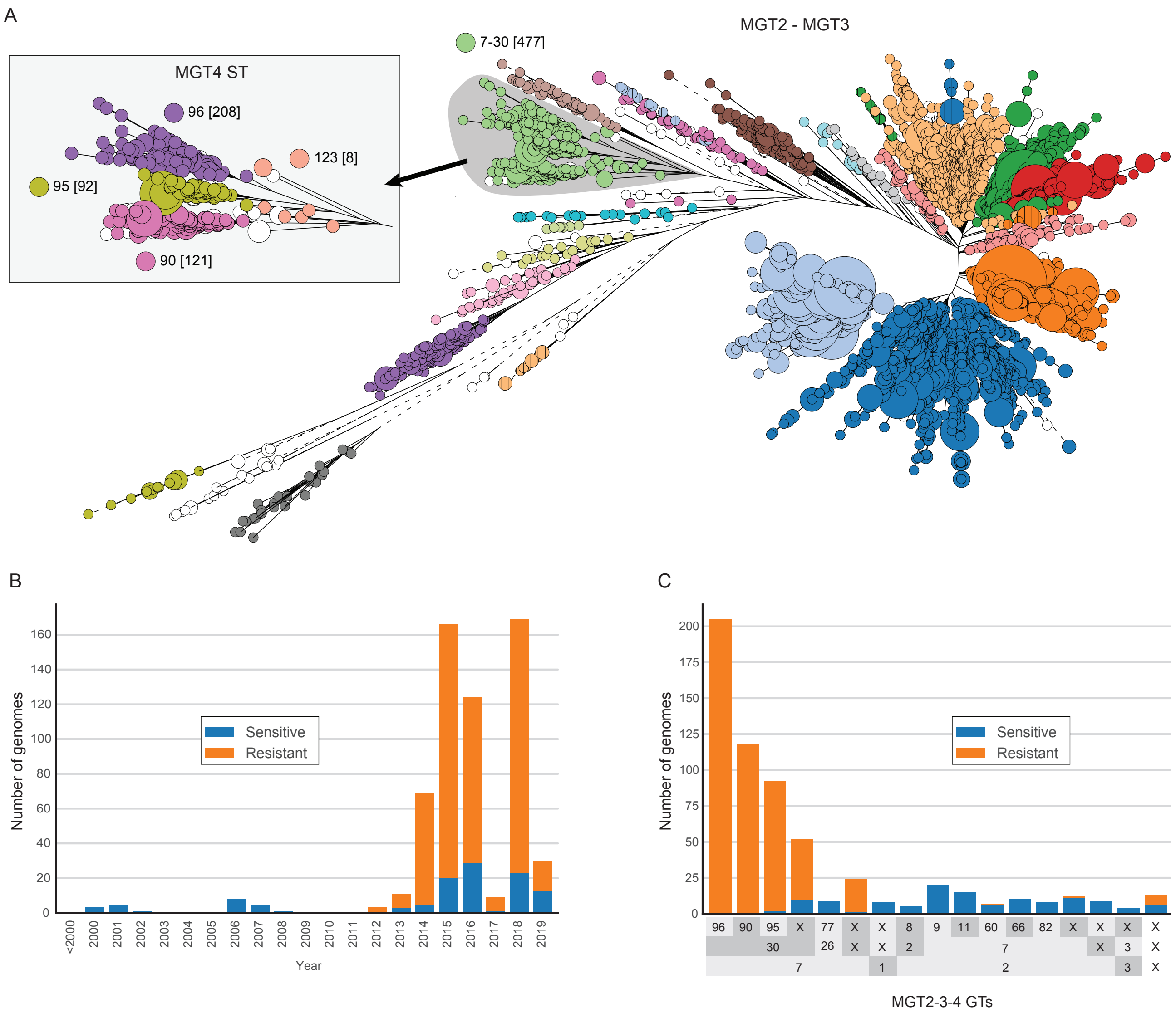


**Supplementary Figure 3. Description of three erythromycin resistant lineages from China. A.** A neighbour joining phylogeny generated from MGT5 allele profiles. Nodes are coloured by MGT2 and MGT3 STs as indicated by labels (identical to figure 2). The inset shows an expanded view of the MGT3 ST30 clade coloured by MGT4 ST. The three resistant lineages are MGT4 ST90, MGT4 ST95 and MGT4 ST96. **B.** All genome sequences available from China up to 2019 grouped by year and coloured by predicted erythromycin resistance. **C.** All genome sequences available from China with assigned MGT2, MGT3 and MGT4 STs on the X-axis. Predicted erythromycin resistance shown by colour. Columns with the same MGT2 and/or MGT3 STs are grouped together. Isolates from STs containing fewer than 6 isolates were collapsed into a single type and labelled X. e.g. 7-X-X isolates were assigned to MGT2 ST7 but their MGT 3 and MGT4 STs contained fewer than 6 isolates.
