## Supplementary Figure 4 for "Genomic epidemiology and multilevel genome typing of *Bordetella pertussis*"

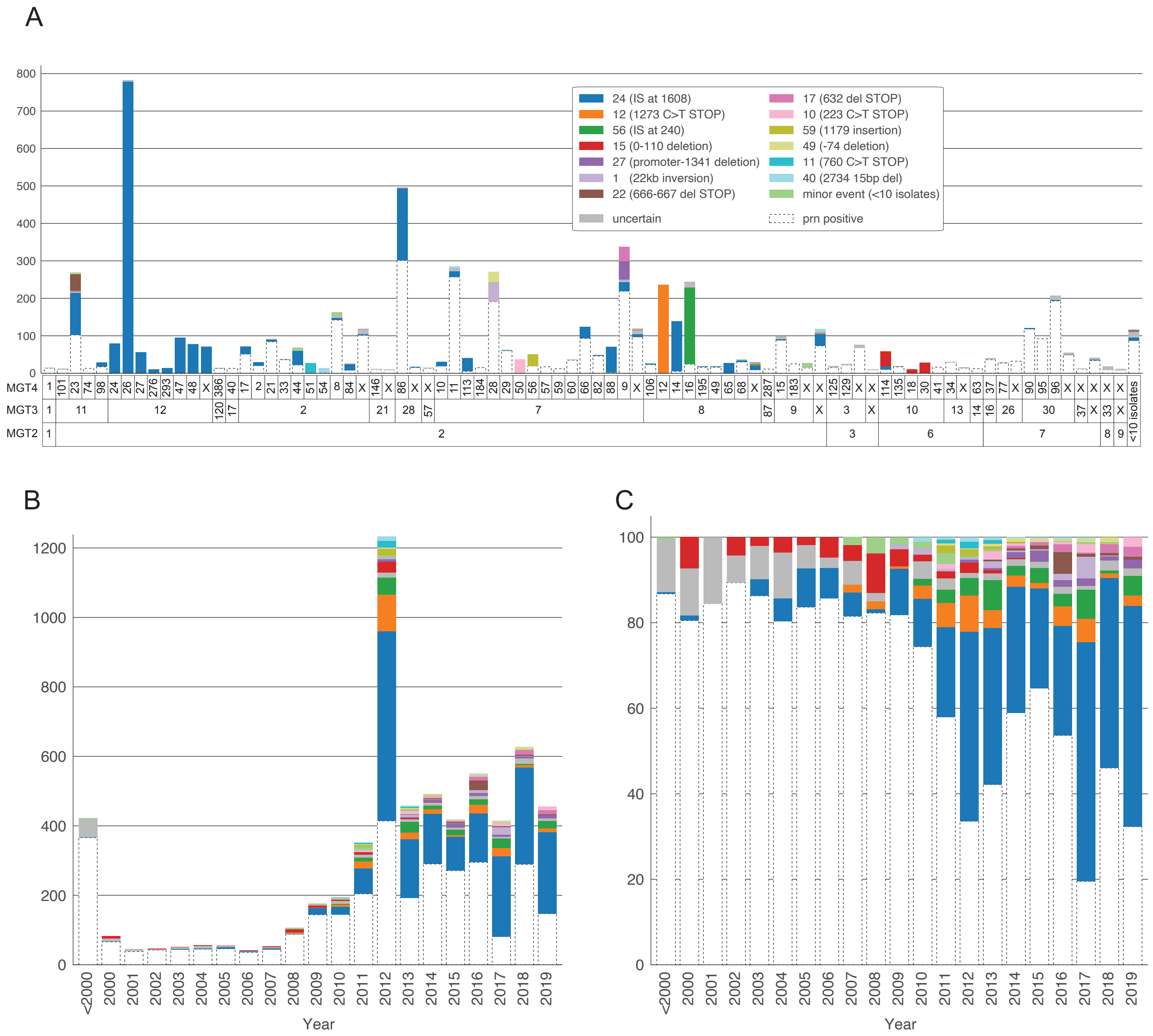


**Supplementary Figure 4. Distributions of prn disruption causes across MGT types and years. A.** Each column is unique set of MGT2, MGT3 and MGT4 STs with 10 or more isolates assigned to it. Each columnis coloured by prn disruption type (colours are the same for all three graphs). prn positive isolates are marker with white dashed outline. The same MGT3 and MGT2 STs are grouped together to allow interpretation at MGT2, MGT3 and MGT4 levels. Isolates from STs containing fewer than 6 isolates were collapsed into a single type and labelled X. e.g. 9-X-X isolates were assigned to MGT2 ST9 but their MGT 3 and MGT4 STs contained fewer than 6 isolates. **B.** The absolute numbers of isolates that are assigned to each disruption type in each year. **C.** The proportion of isolates that are assigned to each disruption type in each year.
