## Supplementary Figure 5 for "Genomic epidemiology and multilevel genome typing of *Bordetella pertussis*"

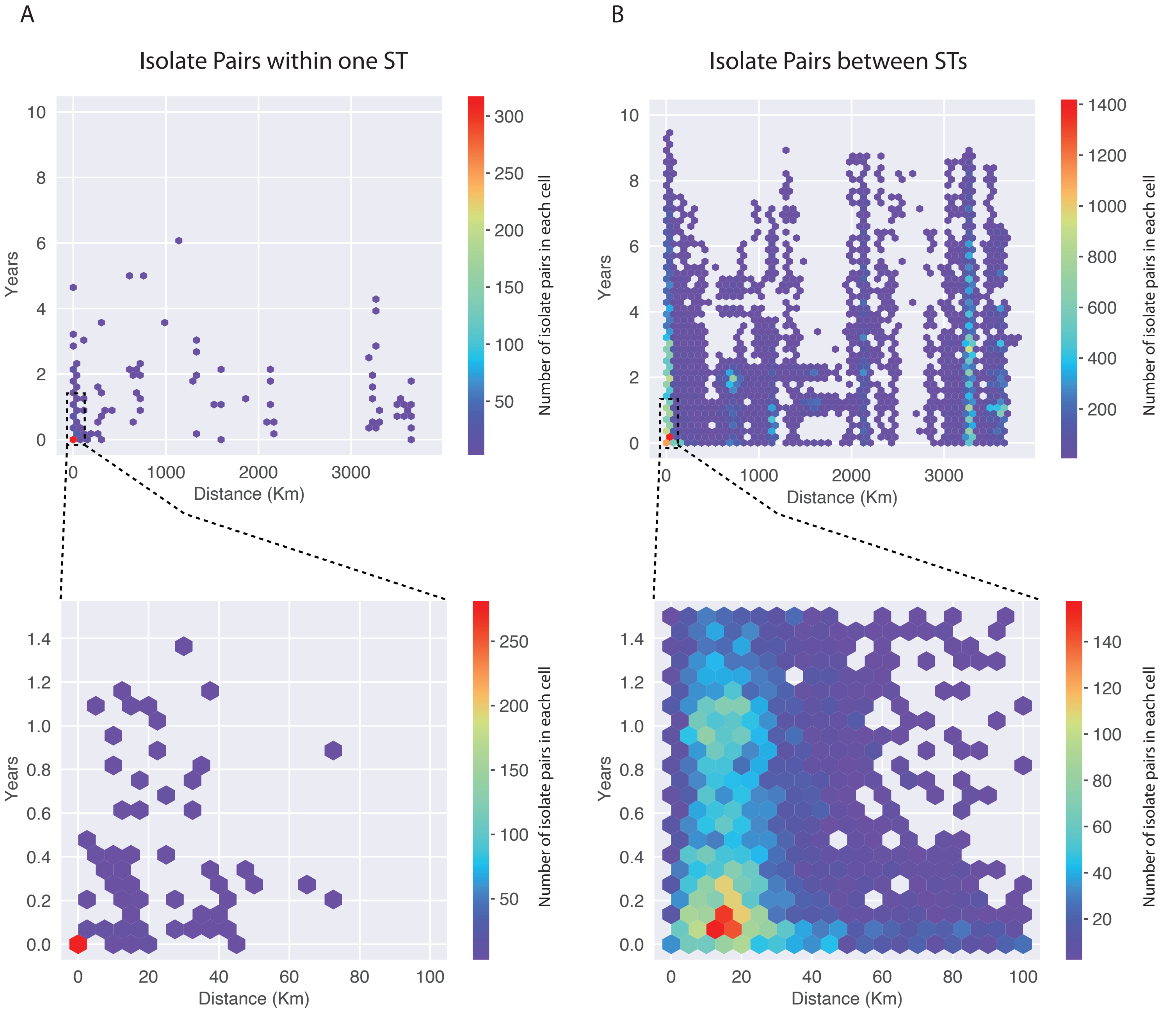


**Supplementary figure 5. Temporal and spatial distributions of isolate pairs within or between MGT5 STs.** Distribution of pairwise distances by year (y axis) and by kilometers (x axis). Data are grouped into cells where the number of isolate pairs falling into each is indicated by its colour. A. Isolate pairs that are both assigned the same MGT5 ST. B. Isolate pairs that are assigned different MGT5 STs. Zoomed in distribution of 0-1.5 years and 0-100km are shown beneith each main plot.
